# Identification of Novel Genes and Variations Associated to Glycolytic Potential Based on Pig Model

**DOI:** 10.1101/367581

**Authors:** Wangjun Wu, Zengkai Zhang, Zhe Chao, Bojiang Li, Caibo Ning, Aiwen Jiang, Chao Dong, Wei Wei, Jie Chen, Honglin Liu

**Affiliations:** Department of Animal Genetics, Breeding and Reproduction, College of Animal Science and Technology, Nanjing Agricultural University, Nanjing, 210095, China; Institute of Animal Sciences & Veterinary, Hainan Academy of Agricultural Sciences, Haikou 571100, China

**Keywords:** pig, transcriptome, glycolytic potential, glycogen, meat quality

## Abstract

In livestock, glycolytic potential (GP) is a critical indicator for evaluating the meat quality. To date, two major genes *protein kinase AMP-activated γ3 non-catalytic subunit gene* (*PRKAG3*) and *phosphorylase kinase catalytic subunit gamma 1*(*PHKG1*), and corresponding cause mutations influencing GP have been confirmed in pigs. Therefore, the aim of this study to identify the novel candidate genes and variations related to GP-related traits using a four-hybrid pig model [Pietrain (P)× Duroc (D)] ×[(Landrace) ×(Yorkshire)]. We totally constructed six RNA-seq libraries using *longissimus dorsi* (LD) muscles, and each library contained two higher GP (H) or two lower GP (L) individuals. A total of 525, 698 and 135 differentially expressed genes (DEGs) were identified between H11 vs L11, H9 vs L9, and H5 vs L5 groups using PossionDis method, respectively. Notably, we found 97 non-redundant DEGs were mapped to GP related QTLs from three paired comparison groups. Moreover, 69 DEGs were identified between H (H11, H9 and H5) and L (L11, L9 and L5) groups using NOIseq method. Additionally, 1,076 potential specific SNPs were figured out between H and L groups, and approximately 40 large Indels with a length ≥ 5bp were identified in each sequencing library. In conclusion, our data provide foundation for further confirming the key genes and the functional mutations affecting GP-related traits in pigs, and also pave the way for elucidating the underling molecular regulatory mechanisms of glycogen metabolism in future study. Moreover, this study might provide valuable information for study on human glycogen storage diseases.

## INTRODUCTION

Pork is the most commonly consumed meat worldwide, its quality have received much attention due to the increase of inferior meat caused by high-strength selection for leaner pork products. However, improving meat quality is one of the most challenging tasks in pig breeding because that it hard to get effective genetic progress using traditional conventional breeding methods. Therefore, identification and application of major genes and variations have become the research hotspots in pig molecular breeding.

Meat is the product of skeletal muscle which is subjected to a serial of structural and biochemical changes after animal slaughter. During the transformation of muscle to meat, the speed and extent of pH value change postmortem are key factors affecting meat quality, such as water-holding capacity (WHC), tenderness, color *etc* (Bee et al. 2007; Popp et al. 2015; Werner et al. 2010). Notably, pH value is directly influenced by glycolysis, with glycogen being converted to lactate and H^+^. Therefore, glycogen, the substrate of glycolysis, its content largely determines the pH value change and closely related to many meat quality traits (Hamilton et al. 2003; Zybert et al. 2013).

The quality of pork is determined by multiple of factors, including genetics, nutrition, feeding methods, slaughtering procedures, and handling of carcass(Jasinska and Kurek 2017; Lebret 2008; Pugliese and Sirtori 2012). Undoubtedly, genetics is the most essential factor affecting meat quality. Currently, *ryanodine receptor 1* (*RYR1*), *protein kinase AMP-activated γ3 non-catalytic subunit gene* (*PRKAG3*; also known as *rendement napole* (*RN*)) are the major genes affecting meat quality. *RYR1*, is a calcium-dependent calcium release channel in the sarcoplasmic reticulum of skeletal muscle, and the g.1843C>T variation in the coding region has been confirmed as the cause mutation for Pale, Soft, and Exudative (PSE) meat (Fujii et al. 1991).

*PRKAG3*, the first major gene affecting glycogen content, its R200Q mutation significantly increases ∼ 70%muscle glycogen content in *RN^-^*animals, which eventually lead to acid meat (Milan et al. 2000). Recently, the second glycogen related major gene, *phosphorylase kinase catalytic subunit gamma 1*(*PHKG1*), its mutation (g.8283C>A) in its ninth intron causes a 32 bp deletion in the open reading frame, generating a premature stop codon, which has been demonstrated to alter GP (43%) and decreases WHC (>20%)(Ma et al. 2014). Notably, the effects of the cause mutations in *PRKAG3 and PHKG1*were observed specifically in Hampshire and Duroc breeds, respectively. In addition, although many different QTLs related to glycogen and related traits have been deposited in pigQTLdb (http://www.animalgenome.org/cgi-bin/QTLdb/SS/index) and some candidate genes and variations related to GP have been investigated (Kaminski et al. 2010; Sieczkowska et al. 2010), the knowledge of major genes and variations related to glycogen and related traits are still limited.

Therefore, the objective of this study was to identify the potential candidate genes related to glycogen metabolism traits using RNA-seq technology with extremely GP phenotypic pigs, and find potential functional variations affecting GP phenotypes through SNP analysis. Our results could provide valuable references for identification of major genes and variations related to glycogen metabolism traits.

## MATERIALS AND METHODS

### Animals and ethics statement

A total of twelve experimental pigs derived from a 279 four-hybrid pigs which crossed by six F_1_ (Pietrain × Duroc) boars and twenty five F_1_ (Landrace× Yorkshire) sows were used for transcriptome analysis. The (P×D)×(L×Y) commercial pig population were reared indoors with uniform environment, and fed with a standard commercial feed ration with free access to water. All the pigs were slaughtered humanely in a standardized commercial abattoir (Jiangsu Sushi Group, China) at an average age of 176 days with an average body weight 110kg in eleven batches (No.1–11) from 1^st^ to 2^rd^ September, 2015. The number of pigs in each batch varied from 23 to 31 pigs. All procedures involving animals were approved by the Institutional Animal Care and Use Committee of Nanjing Agricultural University, Nanjing, Jiangsu, P.R. China (SYXK2011-0036).

### Samples collection and glycolytic potential determination

The LD muscles from all the experimental individuals were excised, and snap frozen in liquid nitrogen at approximately 45min postmortem. Subsequently, the glycolytic potential was determined with 2g muscle tissue between 1th and 2th thoracic vertebrae from posterior. Muscle glucose (MG), glycogen (G), and lactate content were determined using the glucose, glycogen and lactate test kit (Jiancheng, Nanjing), respectively. The glucose-6-phosphate (G6P) was determined using the Glucose-6-Phosphate Assay Kit (Sigma, Spruce Street, St. Louis, MO, USA). Residual glycogen (RG) was the combined content of glycogen and glucose. Glycolytic potential (GP) was calculated according to the formula proposed by Monin and Sellier (Monin and Sellier 1985). GP = 2 × ([glycogen] + [glucose] + [glucose-6-phosphate]) + (lactate). GP was represented as micromoles of lactate equivalent per gram of fresh tissue. In addition, the carcass weight (CW), drip loss measured at 24h postmortem (DL_24h_), drip loss measured at 48h postmortem (DL_48h_), cooking loss (CL), and pH values at 45min and 24h postmortem were measured according the international and national standard protocol. In addition, based on the results of GP phenotypic values, twelve experimental pigs from three slaughtered batches (No. 5, 9 and 11) with the most significant differences between H and L groups were selected for further experiment. A total of twelve economic traits were recorded, and the phenotypic significant differences were statistically analyzed in IBM SPSS 20.0 using unpaired Student’s *t*-test method.

### Total RNA extraction and library construction

The LD muscles from twelve experimental pigs derived from three slaughtered batches (No. 5, 9 and 11), four individuals for each batch (two with the higher GP and two with the lower GP) were selected for construction of transcriptome libraries, and the schematic overview of RNA-seq libraries construction was shown in **Figure S1**. Total RNA of each individual was extracted using the TRIzol reagent (Life Technologies, Carlsbad, CA), separately. The purity and quantity of total RNA were firstly detected using NanoDrop 2000 spectrophotometer (Thermo Scientific), while integrity was assessed using Agilent 2100 (Agilent, Technologies, Inc.). Only the RNA with OD 260/280 between 1.80–2.20 and RIN ≥ 8.0 were used for libraries construction. Briefly, the total RNA from two individuals with the higher GP and two with the lower GP at each slaughtered batch were pooled equally for a sequencing library, respectively. Then, the sequencing libraries were prepared with mRNA-Seq Sample Preparation Kit (Illumina, San Diego, CA, USA) according to the protocol described in the document, and the quality of libraries were validated through checking the size, purity, and the concentration on an Agilent Technologies 2100 Bioanalyzer (Agilent, Technologies, Inc.). Accordingly, a total of six libraries were constructed, and named H11 and L11, H9 and L9, and H5 and L5, separately. ‘H’ and ‘L’ represent the libraries were derived from the higher and the lower GP individuals, respectively.

### Sequencing and raw data processing and alignment analysis

Sequencing was conducted on HiSeq 4000 platform, and 150 bp paired-end reads were produced. The raw data generated were suffered from a serial of filtration, including the adapter, invalid reads containing poly-N and low-quality reads. The remaining high quality reads after filtering were considered as clean reads and saved as FASTQ format. Furthermore, the clean reads were aligned to *Sus scrofa* reference genome (*Sus scrofa* 10.2) using HISAT v0.1.6 (Kim et al. 2015).

### Bioinformatics analysis

Subsequently, transcripts were reconstructed using StringTie software v1.0.4 (Pertea et al. 2015) and novel transcripts were predicted with the genome annotation information using cuffcompare tool (Trapnell et al. 2012), and the differentially splicing genes (DSG) between samples were detected using rMATS v3.0.9 (Shen et al. 2014) and False Discovery Rate (FDR)≤0.05 was set as the threshold value to judge the significance of DSG. After that, the novel coding transcripts were integrated into known transcripts to obtain a complete reference sequence for gene expression analysis. Then, the clean reads to mapped to the reference sequence using Bowtie2 v2.2.5 (Langmead and Salzberg 2012) and expression levels of genes and transcripts were calculated using RSEM v1.2.12 (Li and Dewey 2011).

Furthermore, the differentially expressed genes (DEGs) between the paired samples from the same batch were detected using PossionDis method based on the poisson distribution (Audic and Claverie 1997), and the corrected *P*-value, FDR≤0.001 and absolute value of log2 Foldchange ≥ 1 were considered as DEGs. While, the DEGs between the higher (H) and the lower (L) groups considering biological replicates were detected using NOIseq method based on noisy distribution (Tarazona et al. 2011), and the corrected *P*-value≥0.7 and absolute value of log2 Foldchange ≥ 1 were considered as DEGs.

Thereafter, the DEGs from PossionDis method were mapped to QTL with the pig QTL database (http://www.animalgenome.org/cgi-bin/QTLdb/SS/index) and the QTLs related to glycolytic potential were refined. Additionally, the single nucleotide polymorphisms (SNPs) and insertion-deletion variations (Indels) in each sequencing library were detected with genome mapping result using GATK software v3.4 (McKenna et al. 2010), and the potential differentially and specifically SNPs and Indels were refined manually through comparison analysis between ‘H’ and ‘L’ libraries using the summarized list data of total SNPs **(File S1**). Moreover, the differentially expressed genes containing the potential specifically SNPs were figured out according gene ID and chromosome position of SNPs using SNP annotation information list data (**File S1**) and DEGs annotation information list data (**File S2**).

### Validation of DEGs and SNPs

To validate the reliability of sequencing data, real-time PCR was used to confirm the DEGs. Total RNA were extracted from twelve LD muscle samples for transcriptome sequencing using TRIzol reagent (Invitrogen, Life technologies, Carlsbad, USA), respectively, The first strand cDNA was synthesized using Prime Script^TM^ RT reagent (TaKaRa, Dalian, China), and real-time RT-PCR wasconducted on a Step-One Plus Real-Time PCR System (Applied Biosystems, Carlsbad, CA, USA) using the AceQ qPCR SYBR Green Master Mix (Vazyme, Nanjing, China). A total of nine DEGs were selected for quantitative analysis and all the quantitative primers used in this study were shown in **Table S1**. All reactions were performed in four repetitions and the reference gene *HPRT* was used to normalize gene expression levels. Relative gene expression levels were calculated using the comparative Ct (ΔΔCt) value method. All statistical analyses were performed in IBM SPSS 20.0, and unpaired Student’s *t*-test was used to evaluate the statistical significance of differences. Moreover, correlation analysis was conducted between the fold change values of DEGs from transcriptome sequencing and real-time PCR with Pearson correlation coefficient, *P*<0.05 was considered statistically significant difference. Additionally, eight potential differentially SNPs were validated through PCR product sequencing method, and the information of PCR amplification primers and SNPs location were shown in **Table S2**.

## Data availability

The raw data of transcriptome sequencing data have been deposited to NCBI Short Read Archive (https://www.ncbi.nlm.nih.gov/sra) and are accessible through accession number (SRP127540). **Table S1** contains quantitative primers used in this study. **Table S2** contains sequencing primers and SNPs location information in this study. **Table S3** provides phenotypic data for all experimental pigs with extreme GP value. **Figure S1** is the schematic overview of RNA-seq libraries construction. **Figure S2** depicts the distribution of base quality on clean reads. **File S1** contains the potential differentially and specifically SNPs information identified in this study. **File S2** provides DEGs from PossionDis method. **File S3** contains novel predicted coding_transcripts. **File S4** contains differentially splicing genes. **File S5** provides the genes expression data for all sample libraries. **File S6** provides DEGs from NOIseq method. **File S7** provides the glycolytic potential related QTLs location of DEGs from PossionDis method. **File S8** contains the Indel information identified in this study. **File S9** provides the information on more than 5bp Indel.

## RESULTS

### Phenotypes

In this study, twelve pigs from three batches (four individuals for each batch) represent the higher GP groups (H group: H11, H9, and H5) and the lower GP groups (L group: L11, L9, and L5). H11 and L11, H9 and L9, and H5 and L5 were derived from the same batch, respectively. The phenotypic characteristics of each experimental pig were shown in **Table S3**. From the results, the GP phenotypic values between H and L group individuals were obviously different (**Table 1**). GP phenotypic mean values between H5 and L5, and H9 and L9 were reached extremely significant levels (*P*<0.01). Although the difference of GP phenotypic mean values between H11 and L11 did not reach statistic significant level due to the large variation in H11 group individuals, the obvious difference was actually existed between H11 and L11. In addition, the significant differences of RG, MG, LA, pH_45min_, DL_48h_ and CL between H and L group were also observed.

**Table 1.**
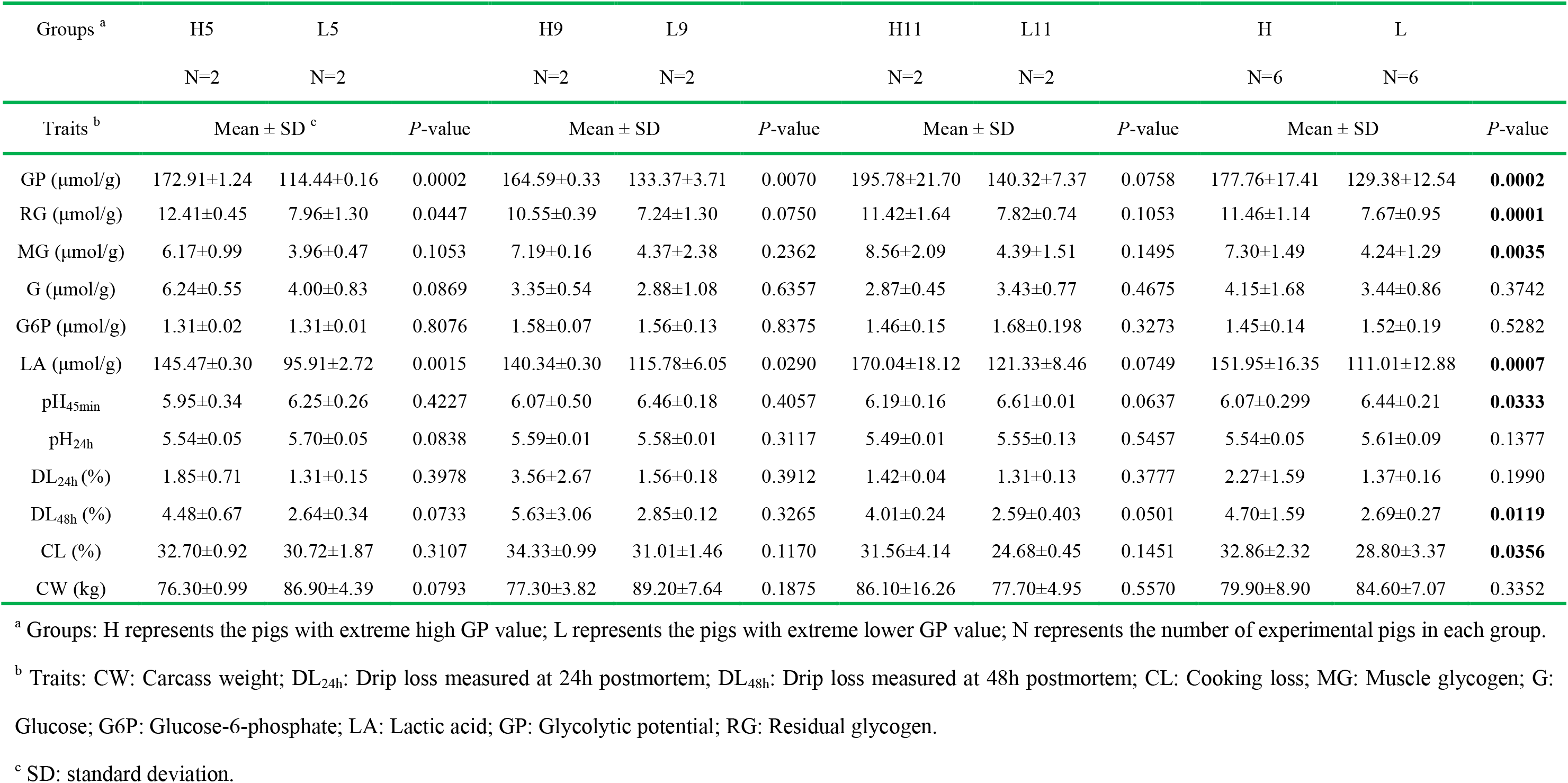
Statistical analysis for experimental pigs with extreme phenotypic GP value

### Sequencing data and alternative splicing analysis

In this study, six sequencing libraries (H11, H9, H5 and L11, L9, L5) were constructed (**Figure S1**) and sequenced on an HiSeq 4000 platform. More than 6 G bases data was generated from each library, representing more than 50.0 million raw reads. The output of data and alignment results were summarized in **Table 2**. After filtering, about 45.0 million clean reads were obtained, the average ratio of clean reads reached 85.0%, and the ratio of clean reads Q20 (those with a base quality > 20 and error rate < 0.01) was higher than 97%, indicating that high-quality clean reads were obtained from each library (**Figure S2**). Moreover, the mapping analysis of clean reads showed that more than 29.0 million clean reads were mapped to reference genome for each library, the ratio reached to 65.0%. In the present study, about 14,000 known genes and hundreds of novel genes were identified in each library, representing approximate 20,000 known transcripts and 7,000 novel transcripts (**Table 2**). The sequence information of novel transcripts can be found in **File S3–1 and File S3–2**. In addition, alternative splicing (AS) analysis showed that five types of AS events were detected, including alternative 3’ splice site (A3SS), alternative 5’ splice site (A5SS), skipping exon (SE), mutually exclusive exon (MXE), and retained intron (RI). The differentially splicing genes (DSG) between samples were shown in **File S4**.

**Table 2.** Output statistics and annotation information of sequencing reads

| Samples | H11 | H9 | H5 | L11 | L9 | L5 |
| --- | --- | --- | --- | --- | --- | --- |
| Total Raw Reads | 52261610 | 53893752 | 52261668 | 52261602 | 53893960 | 50628480 |
| Total Clean Reads | 44993154 | 45347028 | 45385424 | 45136800 | 45134096 | 44466302 |
| Total Clean Reads Ratio (%) | 86.09 | 84.14 | 86.84 | 86.37 | 83.75 | 87.83 |
| Total Adatper Reads | 6210206 | 7439632 | 5912532 | 5982576 | 7761344 | 5190604 |
| Total Adatper Reads Ratio (%) | 11.88 | 13.80 | 11.31 | 11.45 | 14.40 | 10.25 |
| Total High Unknown Base Reads | 21968 | 22364 | 10256 | 21902 | 21254 | 14352 |
| Total High Unknown Base Reads Ratio (%) | 0.04 | 0.04 | 0.02 | 0.04 | 0.04 | 0.03 |
| Total Low Quality Reads | 1036282 | 1084728 | 953456 | 1120324 | 977266 | 957222 |
| Total Low Quality Reads Ratio (%) | 1.98 | 2.01 | 1.82 | 2.14 | 0.18 | 1.89 |
| Clean reads Q20 (%) | 97.73 | 97.71 | 97.72 | 97.64 | 97.66 | 97.63 |
| Clean reads Q30 (%) | 93.62 | 93.58 | 93.85 | 93.44 | 93.43 | 93.6 |
| Total mapped reads | 29629518 | 29244540 | 29843848 | 29824708 | 29420544 | 29634178 |
| Total_mapped_reads ratio (%) | 65.85 | 64.49 | 65.76 | 66.08 | 65.18 | 66.64 |
| Unique_mapped_reads | 26859358 | 26362290 | 26904222 | 26903986 | 26632142 | 26760830 |
| Unique_mapped_reads ratio (%) | 59.70 | 58.13 | 59.28 | 59.61 | 59.01 | 60.18 |
| Multiple mapped reads | 2770160 | 2882250 | 2939626 | 2920722 | 2788402 | 2873348 |
| Multiple mapped reads ratio (%) | 6.16 | 6.36 | 6.48 | 6.47 | 6.18 | 6.46 |
| Splice mapped reads unmapped | 15363636 | 16102488 | 15541576 | 15312092 | 15713552 | 14832124 |
| Splice mapped reads ratio (%) | 34.15 | 35.51 | 34.24 | 33.92 | 34.82 | 33.36 |
| Total Gene Number | 14768 | 13841 | 13976 | 14201 | 14373 | 14058 |
| Known Gene Number | 14067 | 13187 | 13312 | 13516 | 13687 | 13394 |
| Novel Gene Number | 701 | 654 | 664 | 685 | 686 | 664 |
| Total Transcript Number | 20554 | 19069 | 19210 | 19664 | 19959 | 19345 |
| Known Transcript Number | 13385 | 12221 | 12310 | 12600 | 12846 | 12423 |
| Novel Transcript Number | 7169 | 6848 | 6900 | 7064 | 7113 | 6922 |

### Identification of DEGs

Expression levels of genes and transcripts described by Fragments Per Kilobase of transcript per Million fragments mapped (FPKM) in each sample were calculated, the results were shown in **File S5**. On this basis, the DEGs with significance between H11 vs L11, H9 vs L9, and H5 vs L5 were analyzed using PossionDis method. The number of DEGs identified between H11 vs L11, H9 vs L9, and H5 vs L5 were 525, 698 and 135 with FDR≤0.001 and |log2 Foldchange|≥ 1, and the volcanoplots and statistic of DEGs was shown in **Figure 1** and detailed information of DEGs was shown in **File S2**. Moreover, the overlapped DEGs among three groups were analyzed, and 27 DEGs were overlapped between H11 vs L11 (HL11) and H5 vs L5 (HL5), 26 DEGs were overlapped between H9 vs L9 (HL9) and H5 vs L5 (HL5), 157 DEGs were overlapped between H11 vs L11 (HL11) and H9 vs L9 (HL9), while only 6 DEGs were overlapped among three groups (**Figure 2**). In addition, the DEGs between H and L groups considering biological replicates were detected using NOIseq method. A total of 69 DEGs with the corrected *P*-value≥0.7 and |log2 Foldchange|≥ 1 were detected in this study (**File S6**). Moreover, 36 DEGs from NOIseq method were overlapped with that from PossionDis method (**Table 3**). Furthermore, nine DEGs were selected to validate the reliability of sequencing data, and the results indicated that the expression trends of these DEGs were consistent with those obtained from transcriptome data (**Figure 3**), and genes name, genes ID and primers information were shown in **Table S1**. Moreover, correction analysis results showed that the fold change values from the two methods were highly correlated (Pearson correlation coefficient R = 0.87) at a high level of statistical significance (*P*<0.01). These results suggest that reliable sequencing data are obtained from genome-wide transcriptome.

**Figure 1.**
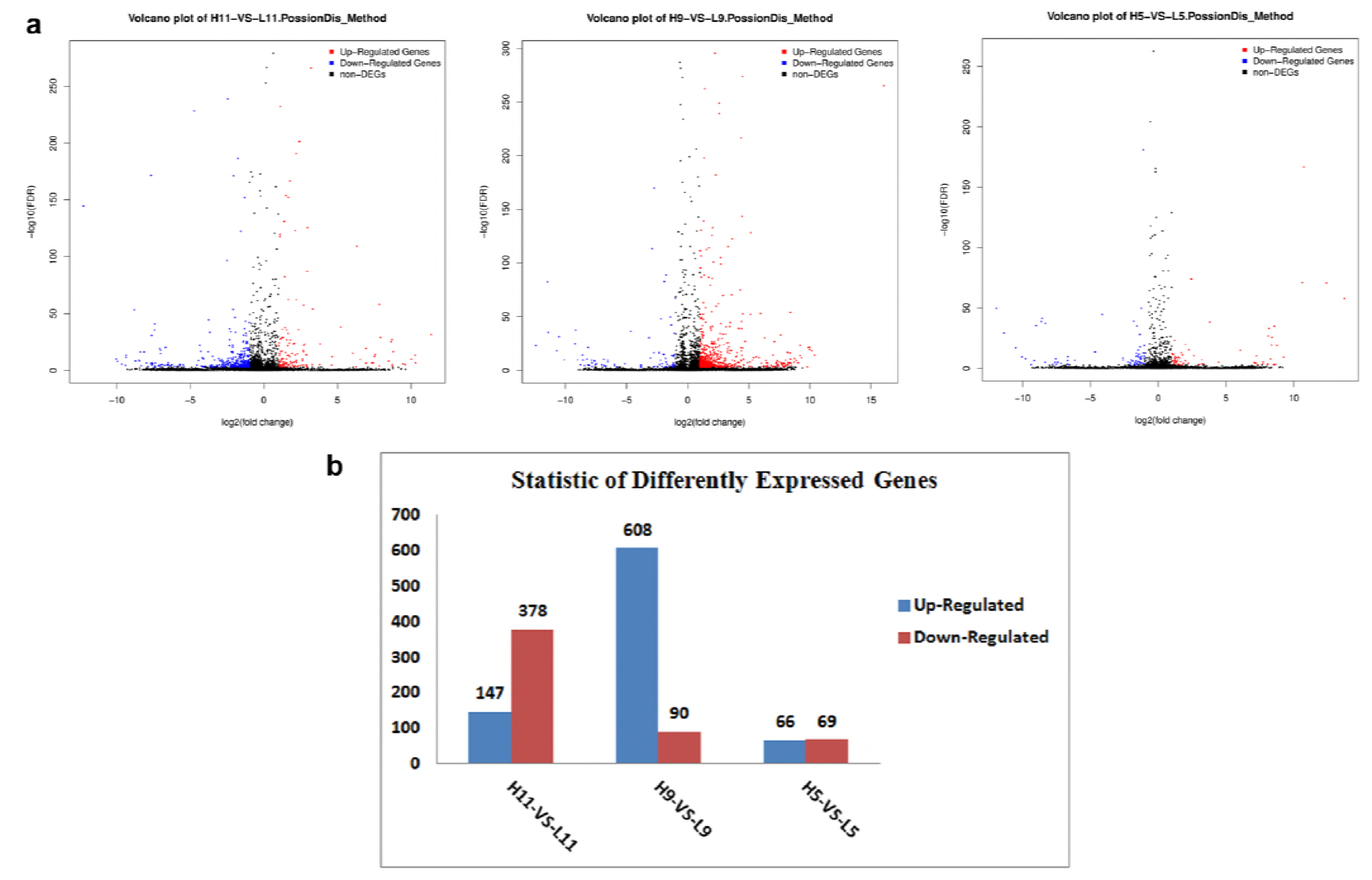
Volcano-plot and statistic of DEGs from PossionDis method. (**a**) Red and blue color represents significantly up-regulated and down-regulated genes, respectively. (**b**) H11-vs-L11, H9-vs-L9 and H5-vs-L5 indicate the comparative strategy by PossionDis method in X axis. Y axis indicates the gene number.

**Figure 2.**
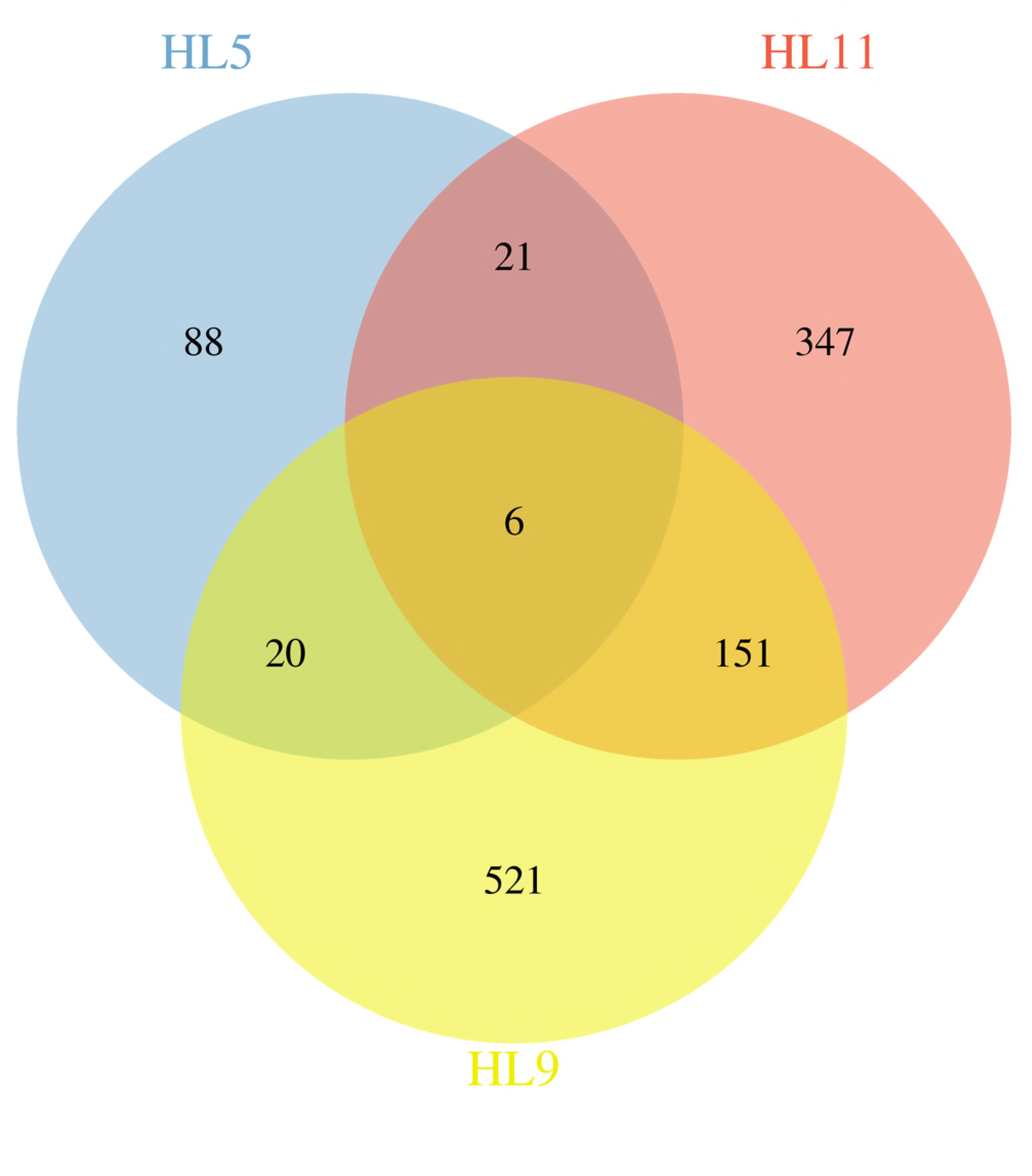
Venn of DEGs obtained from PossionDis method. Overlapping analysis of DEGs was performed using web tool (http://bioinformatics.psb.ugent.be/webtools/Venn/). Different color represents different combination, and the number in the overlap region represents the overlapped DEGs number. In total, 27 DEGs were overlapped between H11 vs L11 (HL11) and H5 vs L5 (HL5), 26 DEGs were overlapped between H9 vs L9 (HL9) and H5 vs L5 (HL5), 157 DEGs were overlapped between H11 vs L11 (HL11) and H9 vs L9 (HL9), and 6 DEGs were overlapped among three groups.

**Table 3.**
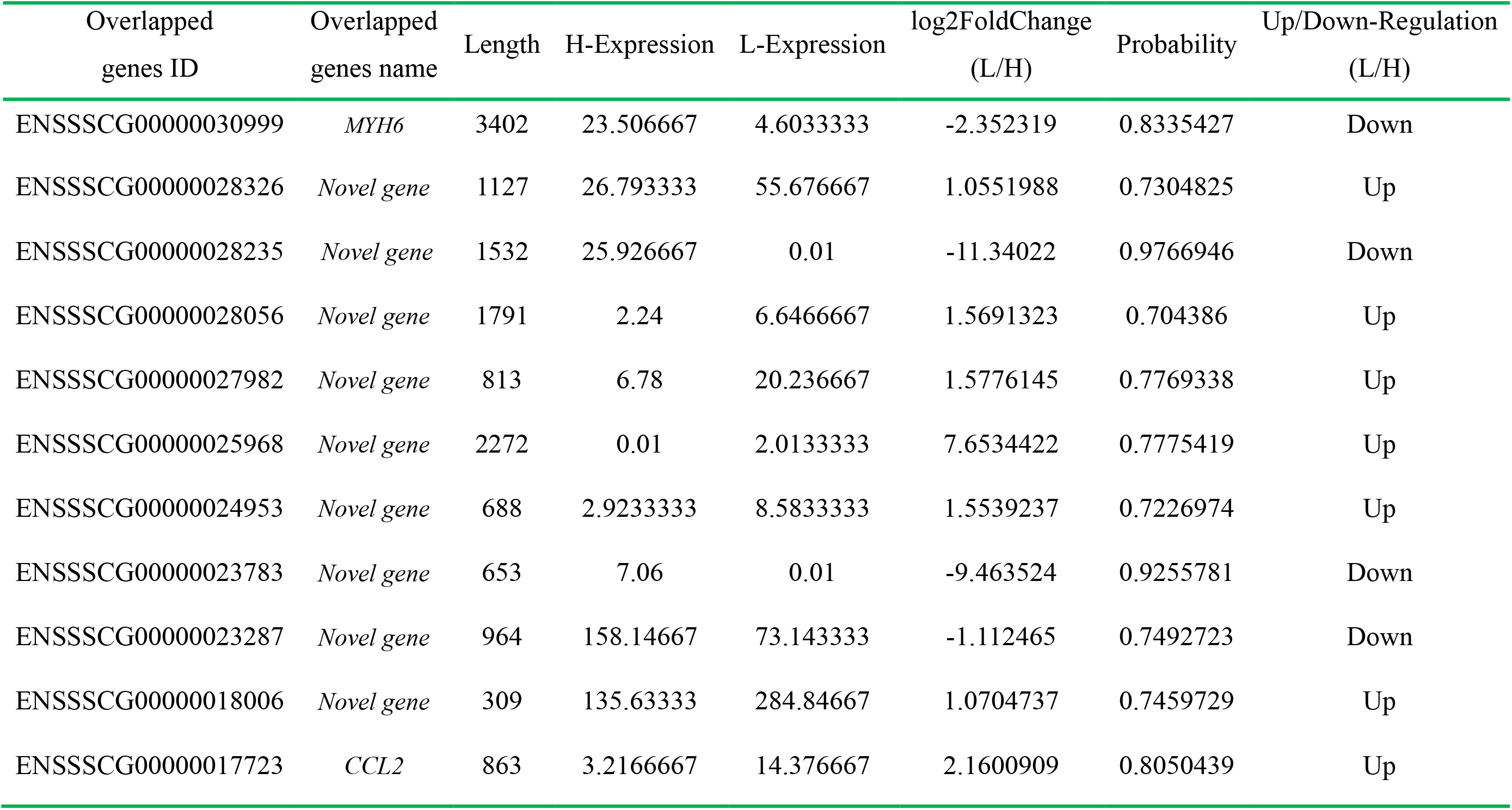
Overlapped DEGs from NOIseq method compared with PossionDis method

| Overlapped<br>genes ID | Overlapped<br>genes name | Length | H-Expression | L-Expression | log2FoldChange<br>(L/H) | Probability | Up/Down-Regulation<br>(L/H) |
| --- | --- | --- | --- | --- | --- | --- | --- |
| ENSSSCG00000030999 | *MYH6* | 3402 | 23.506667 | 4.6033333 | -2.352319 | 0.8335427 | Down |
| ENSSSCG00000028326 | *Novel gene* | 1127 | 26.793333 | 55.676667 | 1.0551988 | 0.7304825 | Up |
| ENSSSCG00000028235 | *Novel gene* | 1532 | 25.926667 | 0.01 | -11.34022 | 0.9766946 | Down |
| ENSSSCG00000028056 | *Novel gene* | 1791 | 2.24 | 6.6466667 | 1.5691323 | 0.704386 | Up |
| ENSSSCG00000027982 | *Novel gene* | 813 | 6.78 | 20.236667 | 1.5776145 | 0.7769338 | Up |
| ENSSSCG00000025968 | *Novel gene* | 2272 | 0.01 | 2.0133333 | 7.6534422 | 0.7775419 | Up |
| ENSSSCG00000024953 | *Novel gene* | 688 | 2.9233333 | 8.5833333 | 1.5539237 | 0.7226974 | Up |
| ENSSSCG00000023783 | *Novel gene* | 653 | 7.06 | 0.01 | -9.463524 | 0.9255781 | Down |
| ENSSSCG00000023287 | *Novel gene* | 964 | 158.14667 | 73.143333 | -1.112465 | 0.7492723 | Down |
| ENSSSCG00000018006 | *Novel gene* | 309 | 135.63333 | 284.84667 | 1.0704737 | 0.7459729 | Up |
| ENSSSCG00000017723 | *CCL2* | 863 | 3.2166667 | 14.376667 | 2.1600909 | 0.8050439 | Up |

**Figure 3.**
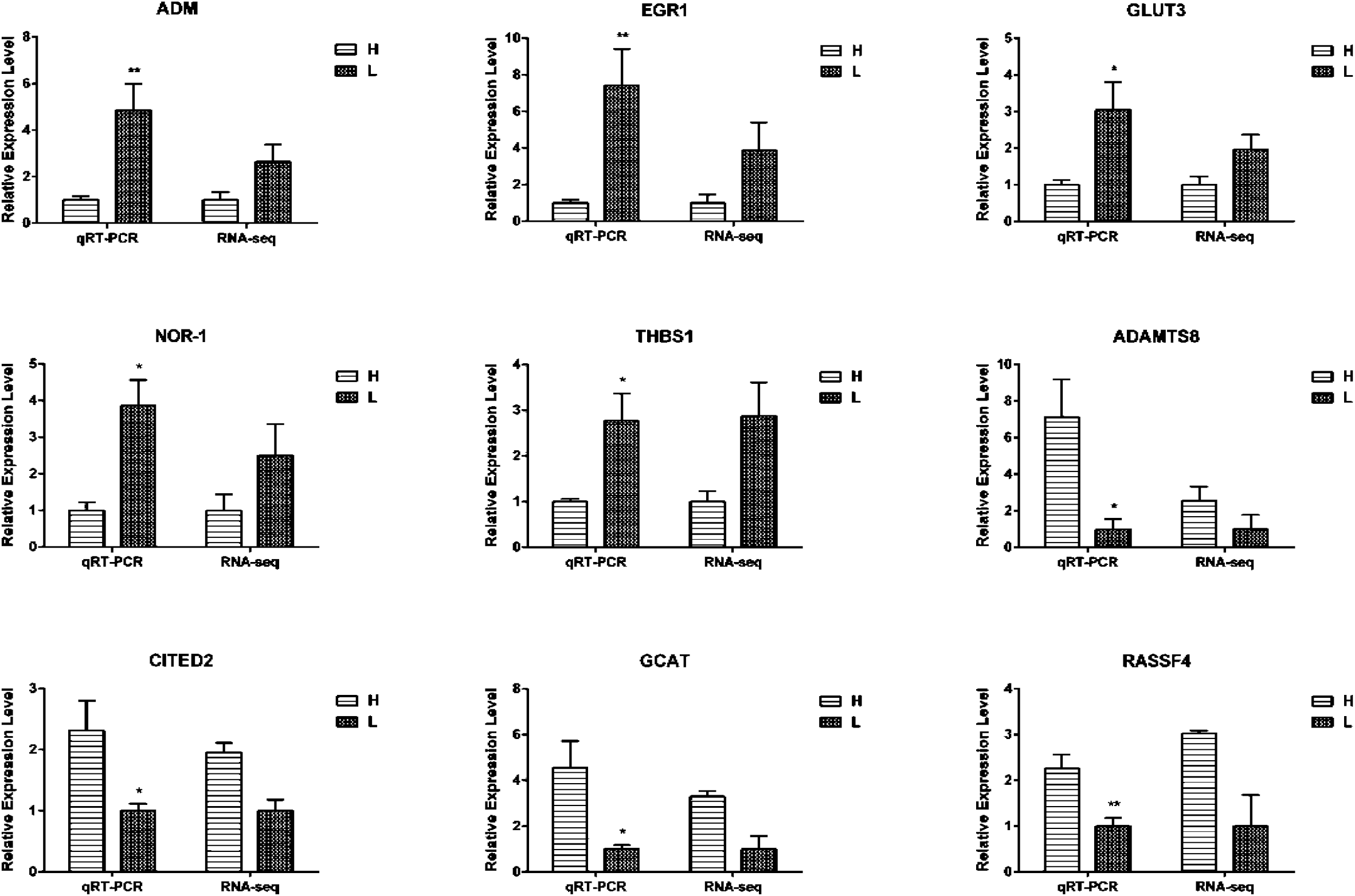
Validation of DEGs by real-time PCR. Total RNA of twelve LD muscle samples were extracted, which derived from three slaughtered batches (No. 5, 9 and 11), four individuals for each batch (two with the higher GP and two with the lower GP). FPKM values are used to calculate the gene expression in RNA-seq and normalize the expression of one group to “1”. In real-time PCR, relative expression levels are calculated using ΔΔCt value method and normalized by reference gene *HPRT*, and similarly normalize the expression of one group to “1”. The paired Student’s *t*-test is used to evaluate the statistical significance of differences between the two groups, * *P* ≤ 0.05，** *P* ≤ 0.01. All data are presented as mean ± standard error (SE).

### QTL location analysis of DEGs

To obtain potential candidate genes affecting GP, we conducted a mapping analysis of DEGs from PossionDis method with the QTL location information of all traits in the pig QTL database. Furthermore, the DEGs located in GP related QTLs, including average glycogen (GLY), average glycolytic potential (GLYPO), average lactate, Glucose-6-phosphate content (GLU6P) and residual glycogen (RESGLY) were refined. In total, 97 non-redundant DEGs were mapped to GP related QTLs from three paired comparisons (H11vs L11, H9 vs L9, H5 vs L5). Moreover, 41 non-redundant DEGs from H11vs L11, 61 non-redundant DEGs from H9 vs L9, and 12 non-redundant DEGs from H5 vs L5 located in GP related QTLs. The detailed QTL mapping information of DEGs from each paired group was shown in **File S7**.

### Identification of potential differential and specific SNPs and Indels

In the present study, six higher GP pigs and six lower GP pigs were used for sequencing analysis. Therefore, another objective of this study was to find potential differential and specific SNPs and Indels between GP higher and lower groups. SNP variation types and numbers were analyzed through comparison with reference genome. About 70,000 SNPs were identified in each sequencing library, and transition is the main SNP variation type (**File S1–1**). The detailed SNP annotation information for each sequencing library, including chromosome, position, gene region, and gene ID were shown in **File S1–2** to **File S1–7**. Subsequently, the potential differentially and specifically SNPs between H and L GP groups were refined. A total of 86,381 potential differentially SNPs (**File S1–8**), and 1,076 potential specific SNPs were figured out (**File S1–9**). Notably, 40 DEGs derived from PossionDis method contained potential specific SNPs were screened (**Table 4**), while only one DEG from NOIseq method, named C-X-C motif chemokine ligand 10 or C-X-C motif chemokine 10 precursor (CXCL10, ENSSSCG00000008977), contained a potential specific SNP (Chromosome 8, Position 75830488). Furthermore, eight potential differentially SNPs were validated using PCR product sequencing method. The results showed that six of eight were real SNPs, the genotype and allele frequencies of each variation in higher and lower GP groups were shown in **Table 5**.

**Table 4.**
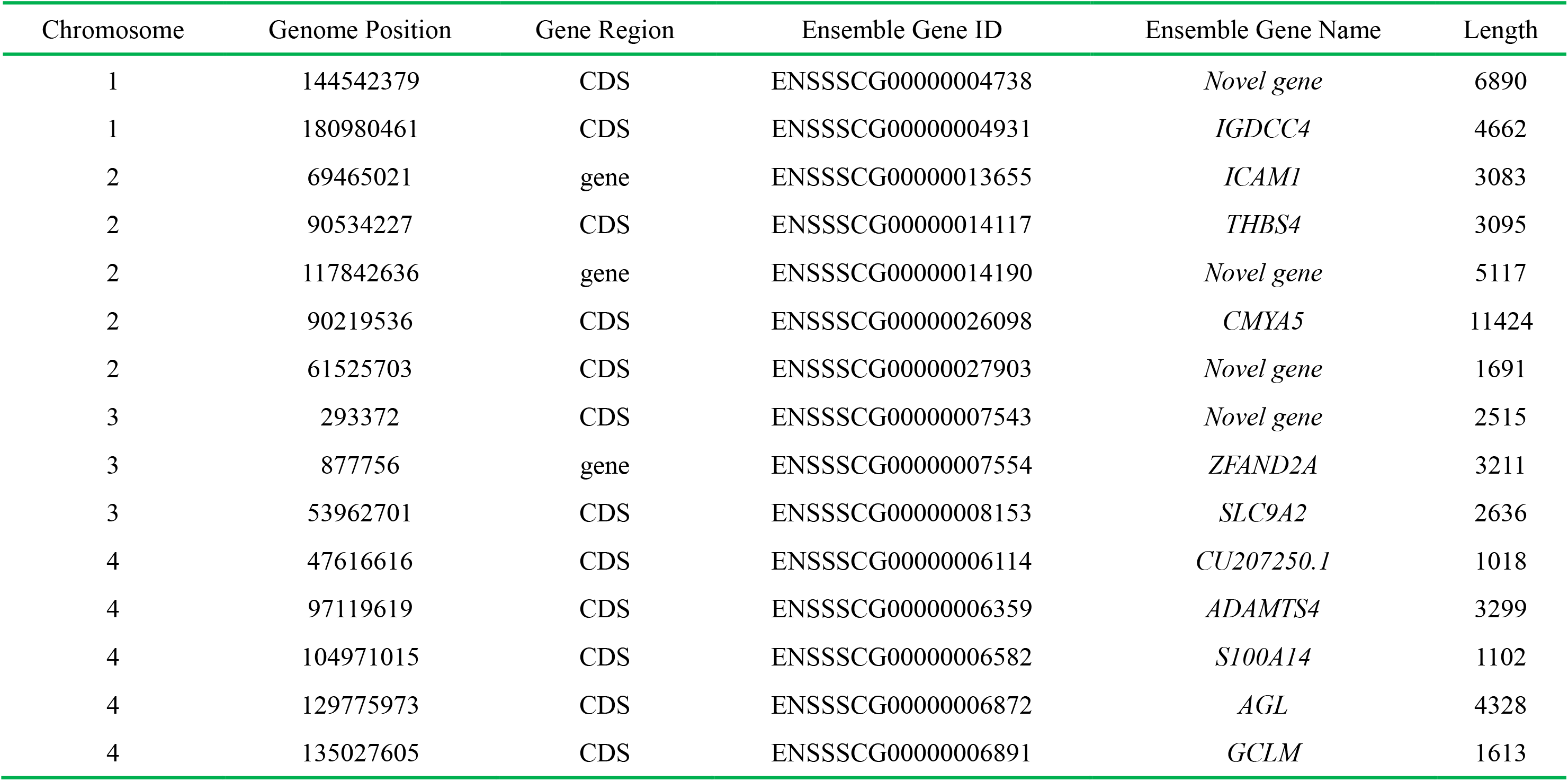
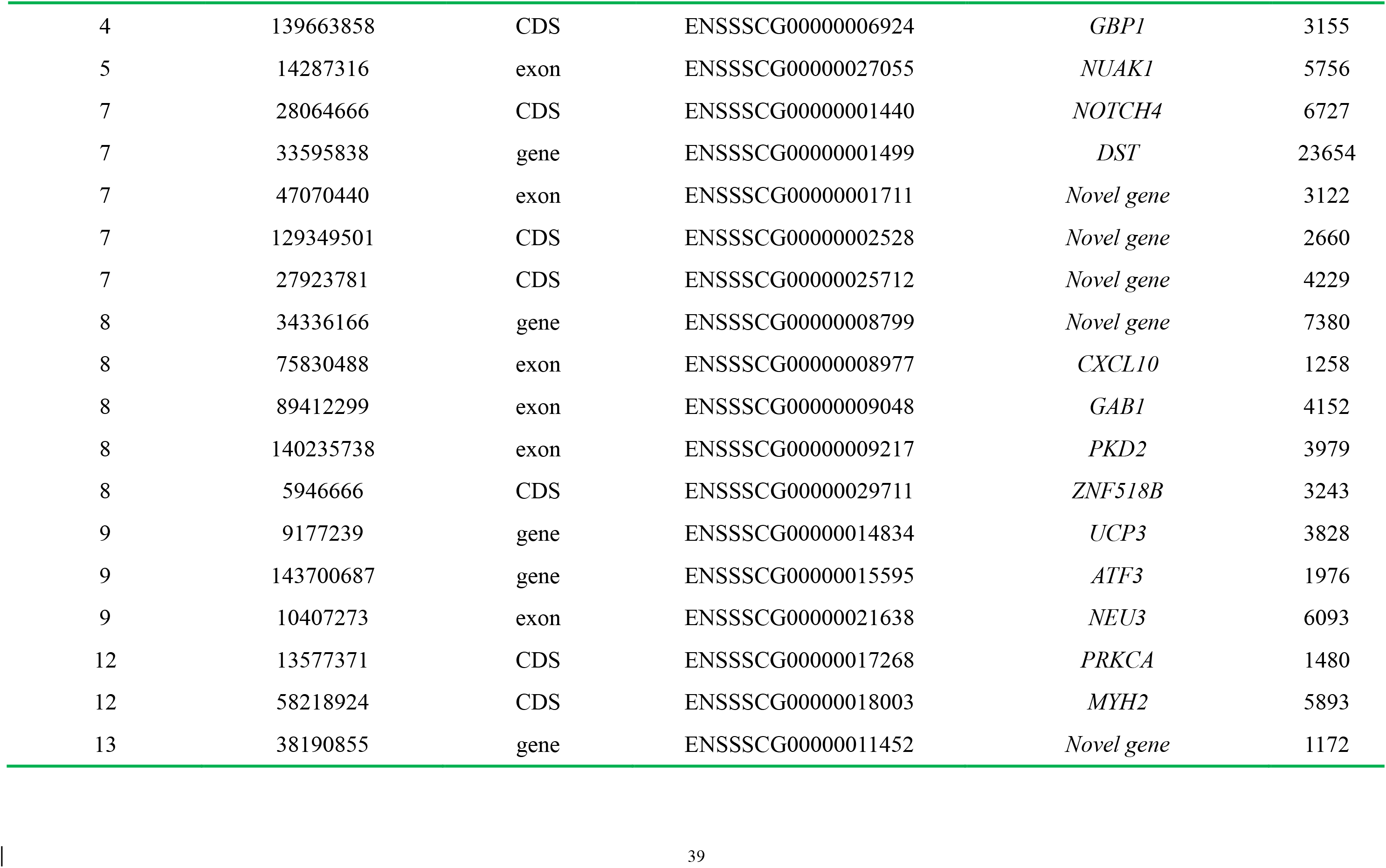
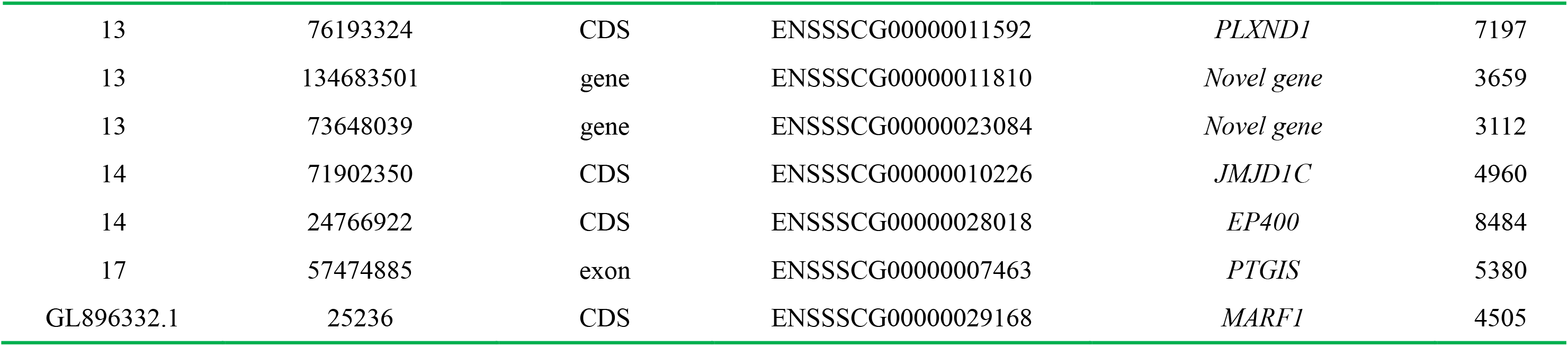
DEGs PossionDis method containing potential specific SNPs

**Table 5.** Validation of the potential differentially SNPs in higher and lower GP groups

|  |  |  |  |  |  |  |
| --- | --- | --- | --- | --- | --- | --- |
| ***AGL gene***<br>(Gene ID: 100156940) |  | Chromosome:<br>4 | Position:<br>129775973 | Variation:<br>T/C |  |  |
|  |  | Genotype |  |  | Allele frequency |  |
|  | Number | C/C | C/T | T/T | C | T |
| H group | 6 | 0 | 0 | 6 | 0.00 | 1.00 |
| L group | 6 | 0 | 4 | 2 | 0.33 | 0.67 |
| ***MAPKB gene***<br>(Gene ID: 100158174) |  | Chromosome:<br>1 | Position:<br>144542379 | Variation:<br>T/C |  |  |
|  |  | Genotype |  |  | Allele frequency |  |
|  | Number | C/C | C/T | T/T | C | T |
| H group | 6 | 1 | 3 | 2 | 0.42 | 0.58 |
| L group | 6 | 3 | 3 | 0 | 0.75 | 0.25 |
| ***CMYA5 gene***<br>(Gene ID: 100505410) |  | Chromosome:<br>2 | Position:<br>90219536 | Variation:<br>T/C |  |  |
|  |  | Genotype |  |  | Allele frequency |  |
|  | Number | C/C | C/T | T/T | C | T |
| H group | 6 | 6 | 0 | 0 | 1.00 | 0.00 |
| L group | 6 | 6 | 0 | 0 | 1.00 | 0.00 |
| ***NEU3 gene***<br>(Gene ID: 100739223) |  | Chromosome:<br>9 | Position:<br>10407273 | Variation:<br>T/C |  |  |
|  |  | Genotype |  |  | Allele frequency |  |
|  | Number | C/C | C/T | T/T | C | T |
| H group | 6 | 0 | 0 | 6 | 0.00 | 1.00 |
| L group | 6 | 0 | 0 | 6 | 0.00 | 1.00 |
| ***NUAK1 gene***<br>(Gene ID: 100523669) |  | Chromosome:<br>5 | Position:<br>14287316 | Variation:<br>T/C |  |  |
|  |  | Genotype |  |  | Allele frequency |  |
|  | Number | C/C | C/T | T/T | C | T |
| H group | 6 | 1 | 4 | 1 | 0.50 | 0.50 |
| L group | 6 | 1 | 3 | 2 | 0.42 | 0.58 |
| ***PRKCA gene***<br>(Gene ID: 397184) |  | Chromosome:<br>12 | Position:<br>13577371 | Variation:<br>T/C |  |  |
|  |  | Genotype |  |  | Allele frequency |  |
|  | Number | C/C | C/T | T/T | C | T |
| H group | 6 | 5 | 1 | 0 | 0.92 | 0.08 |
| L group | 6 | 3 | 3 | 0 | 0.75 | 0.25 |
| ***PTGIS gene***<br>(Gene ID: 100126284) |  | Chromosome:<br>17 | Position:<br>57474885 | Variation:<br>T/C |  |  |
|  |  | Genotype |  |  | Allele frequency |  |
|  | Number | C/C | C/T | T/T | C | T |
| H group | 6 | 2 | 4 | 0 | 0.67 | 0.33 |
| L group | 6 | 5 | 1 | 0 | 0.92 | 0.08 |
| ***UCP3 gene***<br>(Gene ID: 397116) |  | Chromosome:<br>9 | Position:<br>9177239 | Variation:<br>T/C |  |  |
|  |  | Genotype |  |  | Allele frequency |  |
|  | Number | C/C | C/T | T/T | C | T |
| H group | 6 | 0 | 2 | 4 | 0.83 | 0.17 |
| L group | 6 | 1 | 4 | 1 | 0.50 | 0.50 |

Additionally, about 5,000–7,000 Indels were identified from each sequencing library when compared with reference genome (**File S8–1 to File S8–6**), and 292,162 and 66 potential differential Indels were figured out from H11 vs L11 (**File S8–7)**, H9 vs L9 (**File S8–8)**, and H5 vs L5 (**File S8–9**) groups, respectively. Moreover, the number of Indels was reduced as the length increase, **File S9** showed the annotation information of more than 5bp length indels.

## DISCUSSION

Glycogen metabolism disorder involved in defects in glycogen metabolism or excess glycogen stored in liver and muscle cause glycogen storage diseases. In livestock, measurement of muscle glycogen (G) combined with measure of glucose (MG), glucose-6-phosphate (G6P) and lactate content called glycolytic potential (GP) to evaluate the capability of glycogen storage and using it to evaluate the meat quality development. Therefore, we measured the GP phenotypic values of each pig derived from a 279 P×D×L×Y commercial pigs and selected the pigs with extreme GP phenotypic values for RNA-seq analysis in this study. In order to meet the requirements of biological replicates and use as many experimental pigs as possible in sequencing, we selected twelve experimental pigs from three slaughtered batches (No.11, 9 and 5) and four individuals with extreme GP phenotypic values for each batch (two with the higher GP and two with the lower GP) to construct RNA sequencing libraries (**Figure S1**). Factually, the phenotypic values of many other meat quality traits showed obvious differences between the higher and lower GP groups, and several reached statistical significant differences, including pH_45min_, DL_48h_, and CL (**Table 1**). Previous studies indicated that GP was closely related to pH value, drip loss and cooking loss (Hamilton et al. 2003; Zybert et al. 2013), which suggest that the pigs selected are suitable for sequencing analysis.

In the present study, six sequencing libraries were constructed and high-quality sequencing data were obtained through quality-control analysis, including Q20 and Q30, reads mapping, and base composition analysis. The ratio of clean reads with> Q20 and >Q30 reached 97.0% and 93.0% for each library, respectively. The ratio of mapped reads reached more than 65.0% for each library, which is consisted with previous studies (Chen et al. 2011; Esteve-Codina et al. 2011; Li et al. 2016b), but less than the some studies where more than 70.0% of reads were mappable (Li et al. 2016a; Ramayo-Caldas et al. 2012). These discrepancies may be caused by the differences of reference genome version and alignment software. Generally, the reads obtained from six libraries can meet the subsequent analysis of differentially expressed genes identification and variations detection.

One of the main objectives of this study to identify the candidate genes affecting GP related traits. Based on the strategy of RNA-seq libraries construction (**Figure S1**), we adopted two methods to screen the DEGs in the present study. PossionDis method was used to identify the DEGs considering paired comparisons of H11 vs L11, H9 vs L9, and H5 vs L5. Satisfactorily, hundreds of DEGs were found in paired comparative analysis and a certain number of DEGs were overlapped between two groups, although only six DEGs were overlapped among three groups (**Figure 2**). In RNA-seq analysis, some questions were still difficulty to avoid in livestock, such as the large variations were often happened between samples. These phenomenon was also observed in this study, which may be the reason that only a few overlapping genes were identified among each RNA-sequencing libraries. Moreover, the FDR≤0.001 and log2 Foldchange ≥ 1 were considered to screen the DEGs in this study, so some truly DEGs may be ignored. Moreover, NOIseq method was applied to identify the DEGs between H and L groups considering biological replicates, 69 DEGs were detected in this study. Notably, 36 DEGs from NOIseq method were overlapped that from PossionDis method (**Table 3**). Although the major genes related to GP have not been confirmed, the DEGs found in this study pave good foundation for identifying the genuine major genes affecting GP.

At present, some QTLs related to average glycogen (18), average glycolytic potential (20), average lactate (10), glucose-6-phosphate content (3) and residual glycogen (6) were deposited in pig QTLs database, but it is still difficult to confirm the functional genes affecting glycogen metabolism due to the large size of these QTLs. To refine the candidate genes related to glycogen metabolism, we performed an integrated analysis between porcine QTLs and DEGs according the genomics location information. Fortunately, we found 97 potential candidate genes affecting GP-related traits (**File S7**), although the functions of these candidate genes are still needed to confirm. GP respond to the glycogen storage capability, which can be used to evaluate the meat quality development, and is closely related to several meat quality traits. Up to date, only two major genes and corresponding cause mutations affecting GP have been confirmed, one is *PRKAG3*(Milan et al. 2000) and another is *PHKG1*(Ma et al. 2014). Notably, harmful allele (*RN*^-^ allele) in *PRKAG3* gene is exclusively found purebred Hampshire breed and its related synthetic lines, and the effect is only observed in purebred Hampshire breed, which limited its application in most pig breeding programs. *PHKG1* is the second major gene related to glycogen metabolism and g.8283C>A is the cause mutation, while the mutation is mainly observed in Duroc breeds, especially in White Duroc breed (Ma et al. 2014), which can be implicated for screening of excellent Duroc pigs, but the application value in other pig breeds are limited. Therefore, additional major genes and mutations are needed to be explored for GP-related traits molecular breeding in most of pig breeds. In the present study, the experimental pigs are from a four-hybrid commercial pig (P×D) × (L×Y) enriched in genetic diversities, which suggest the candidate genes and variations identified in this study might be applied more widely.

Recent years, the genome sequences and high-density SNP chips are widely used for detection of variations significantly associated to economic traits in livestock (Liu et al. 2015; Ma et al. 2014; Rubin et al. 2012; Rubin et al. 2010; Verardo et al. 2016). In this study, we found 1,076 potential specific SNPs between the higher and the lower GP group (**File S1–9**) using the trancriptome sequencing data, which are the potential variations associated to GP-related traits. Notably, 40 DEGs contains potential specific SNPs (**Table 4**), and most of SNPs are located in gene coding sequence (CDS) which is in accordance with the data from trancriptome sequencing. Factually, many causal mutations of major genes identified in previous studies are in CDS region and cause the change of amino acid, such as theg.1843C>T mutation in *RYR1* gene (Fujii et al. 1991), the R200Q mutation in *PRKAG3* gene (Milan et al. 2000), the G>A mutation in CDS of MC4R gene (Kim et al. 2000). Therefore, the potential specific SNPs identified in this study can provide valuable references for identifying the functional mutations associated with GP-related traits. In the present study, a potential specific SNP was included in *CXCL10*gene, while previous study reported that *CXCL10*was closely related to Type1 diabetes (Antonelli et al. 2014). Intriguingly, a SNP was validated in exon of differentially expressed gene *AGL* in this study. AGL, a key glycogen debrancing enzyme, plays an important role in reducing glycogen degradation and accumulating of limit dextrin in skeletal muscle. It was previously reported that a causal mutation in human AGL exon has close relationship with glycogen storage disease type III (Rousseau-Nepton et al. 2015).

Moreover, the indels were analyzed and hundreds of potential differentially Indels were figured out between the higher and lower GP groups. To our special attention, the effects of indels are often enormous, for example, an 11bp deletion in the third exon of *myostain* gene (*GDF-8*) causes the double-muscled phenotype in Belgian Blue cattle (Grobet et al. 1997; McPherron and Lee 1997). Therefore, although only a few number of large insertion and deletion are observed in our data, approximately 40 Indels with a length ≥ 5bp were identified in each sequencing library (**File S8**), however, the relationship between these indels and GP-related traits, and regulatory mechanisms might become the focus in future study.

## CONCLUSIONS

In livestock, GP, responding to the glycogen storage capability, is a critical indicator for evaluating meat quality. In the present study, we conducted high-throughput trancriptome sequencing to identify the potential candidate genes and variations related to GP-related traits using the muscle samples from a four-hybrid commercial pig (P×D×L×Y). A total of 525, 698 and 135 DEGs were identified between H11 vs L11, H9 vs L9, and H5 vs L5 GP groupsusing PossionDis method, respectively. While, a total of 69 DEGs were identified between H (H11, H9 and H5) and L (L11, L9 and L5) GP groups using NOIseq method. Moreover, 1,076 potential specific SNPs were figured out between H and L GP groups, and approximately 40 large Indels with a length ≥ 5bp were identified in each sequencing library. Taken together, our data provide foundation for further identifying the key genes and screening the functional mutations affecting GP-related traits.

## ACKNOWLEDGEMENTS

We thank Jiaolong Li from Nanjing Agricultural University for his help in samples collection. This work was supported by the National Natural Science Foundation of China (31501920), the Fundamental Research Funds of the Central Universities (KJQN201605), the National Major Project of Breeding for Transgenic Pig (2016ZX08006001-003) and the Program of Hainan Provincial Engineering Research Center for Livestock and Poultry Cultivation (XQ2016JSZX001).

---

**Figure S1.**
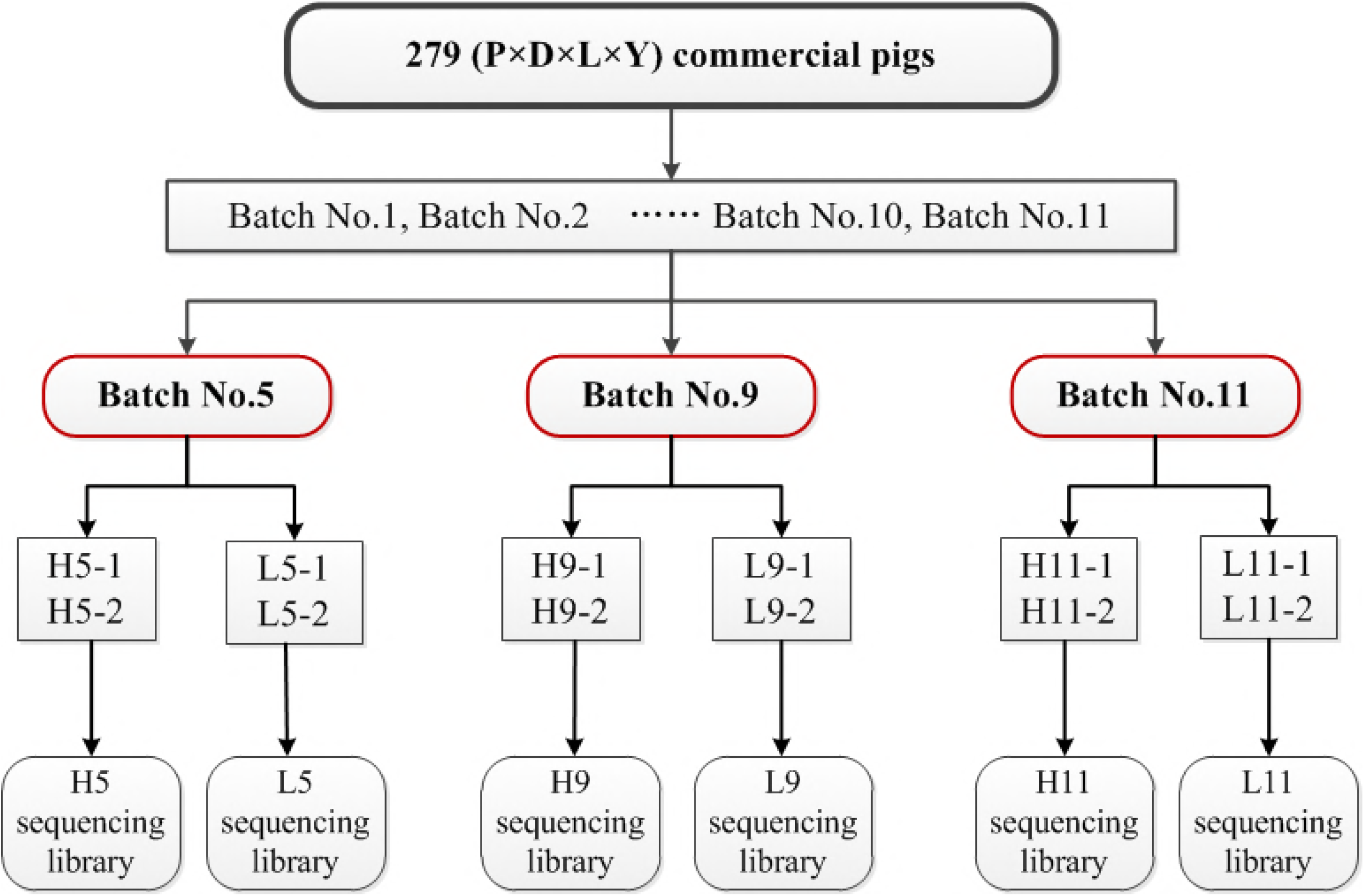
The schematic overview of RNA-seq libraries construction.

**Figure S2.**
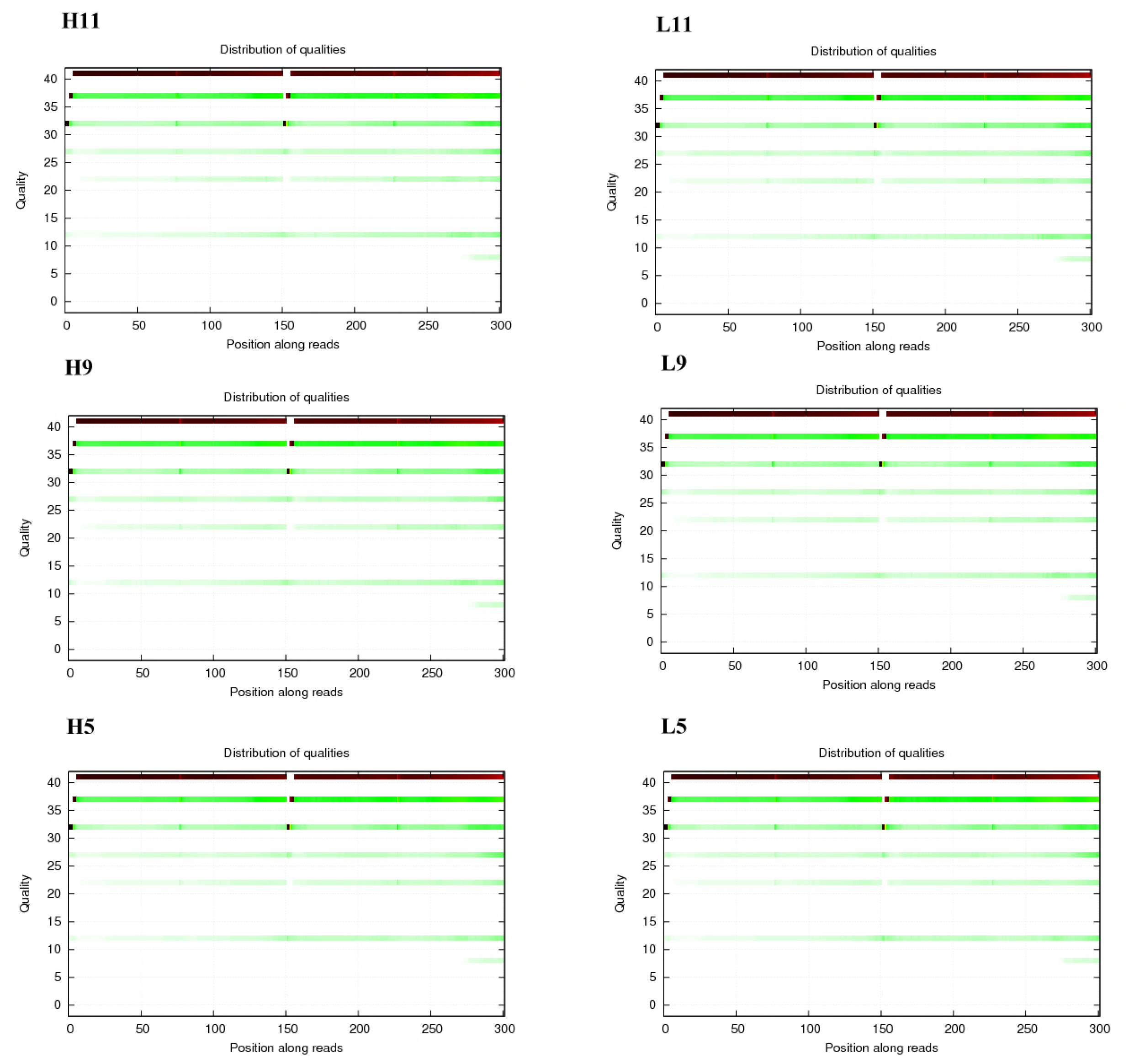
Length distribution of clean reads. Y axis represents the number of total clean reads with certain length. X axis represents the length of clean reads

